# Attention and the motion after effect: New measurements with visual search reveal important individual differences in adaptability

**DOI:** 10.1101/478941

**Authors:** Michael J. Morgan, Joshua A. Solomon

**Affiliations:** Centre for Applied Visual Science, City, University of London

## Abstract

It is usually assumed that sensory adaptation is a universal property of human vision. However, in two experiments designed to measure adaptation without bias, we have discovered a minority of participants who were unusual in the extent of their adaptation to motion. One experiment was designed so that targets would be invisible without adaptation; the other, so that adaptation would interfere with target detection. In the first, participants adapted to a spatial array of moving Gabor patches. On each trial the adapting array was followed by a test array in which but all of the test patches except one were identical to their spatially corresponding adaptors; the target moved in the opposite direction to its adaptor. Participants were required to identify the location of the changed target with a mouse click. The ability to do so increased with the number of adapting trials. Neither search speed nor accuracy was affected by an attentionally-demanding conjunction task at the fixation point during adaptation, suggesting low-level (pre-attentive) sites in the visual pathway for the adaptation. However, a minority of participants found the task virtually impossible. In the second experiment the same participants were required to identify the one element in the test array that was slowly moving: reaction times in this case were elevated following adaptation. The putatively weak adapters from the first experiment found this task easier than the strong adapters.

## Introduction

Our experiment was designed to develop a performance-based (i.e. “Type 1,” Sperling, Dosher, & Landy, 1990) measure of motion adaptation. Performance-based measures have many applications, but we specifically wanted to learn whether the motion aftereffect (MAE) is reduced when observers’ attention is distracted away form the adapting stimulus, as reported by Chaudhuri (1990), Rees, Frith, and Lavie (1997), and Taya, Adams, Graf, and Lavie (2009). This possibility is theoretically interesting because there is good evidence that the locus of adaptation underlying the MAE is visual area V1 (Kohn & Movshon, 2003). An effect of attentional distraction on the MAE would thus imply a top-down effect of attention on visual processing in V1 or earlier.

To date, almost all prior investigations of this question used phenomenological (“Type 2”) measures of the MAE. Prolonged inspection of a moving image such as a waterfall causes subsequently viewed, stationary stimuli to appear as if they are moving in the opposite direction (Addams, 1834; Wohlgemuth, 1911; Mather, Verstraten, & Anstis, 1998). The measurement of this effect by its duration is potentially subject to criterion and expectation effects (Sinha, 1952). It is difficult to decide when a stimulus has stopped moving, particularly if it is known not to be moving in the first place.

Wohlgemuth (1911; see Wade, Thompson, & Morgan, 2014) measured the MAE by its apparent duration. He employed a task involving random, serial, visual presentations (RSVP) that was designed to distract observers from the adaptor. This task did not affect MAE duration in Wohlgemuth’s study. Chaudhuri (1990) and Rees *et al.* (1997), on the other hand, reported positive results with similar methods. A positive result was also reported by Taya *et al.* (2009), who measured the MAE by the speed of test motion required to compensate for (or “null”) it, which has the same problems as the measurement of duration. Nishida & Ashida (2000) found no effect of distraction on MAE duration, but did they did find an effect of distraction on the contrast assigned to a moving test pattern that was required to null the interocular MAE only (i.e. there was no effect on the monocular MAE). Morgan (2011, 2012, 2013) has repeatedly found no effect of distraction on the MAE. The 2012 study used naïve students and MAE duration. It also used more experienced obsevers and the nulling paradigm. The 2013 study measured the MAE by its effect on perceived speed (Thompson, 1981), interleaving various pedestals to defeat any simple decision strategy (e.g. “when in doubt respond that the adapted stimulus is slower”). However, even the 2013 experiment is in essence a Type 2 measure. The 2011 study is the only one to date to use a Type 1 measure, namely the direction-specific loss of contrast sensitivity following adaptation. However, the measurement of contrast sensitvity is lengthy and not well suited to measuring the growth of adaptation over time, and consequently may obliterate attentional effects during the build-up of adaptation. Bartlett, Taya, and Graf (2016) have suggested that attentional distraction might affect the rate of growth of adaptation to asymptote without affecting the asymptote itself. The purpose of the experiments we report here was to develop a rapid, simple Type 1 measure of adaptation that could track the growth of adaptation over trials.

Our new Method is based on visual search (Wissig, Patterson, & Kohn, 2013). Observers adapted to an array of Gabor patches, some of which moved upwards and some downwards (Fig. 1). Periods of adaptation alternated with the presention of a brief test stimulus, with a 1-s temporal gap between adaptor and test to prevent transient-based detection. The test array was identical to the adapt array except that one of the patches (the “target”) reversed its direction from the adapting direction (the “Target-change” condition). All the other patches (the “distractors”) moved in the same direction as their spatially corresponding adaptors. Morgan & Hauperich (2017) reported that in these circumstances the target can pop out from the distractors, and its position can be detected. In the present series of experiments we measure the growth of performance with the duration of prior adaptaton, and the effects of distracting attention from the adapting stimulus with a difficult central RSVP task.

**Fig. 1.**
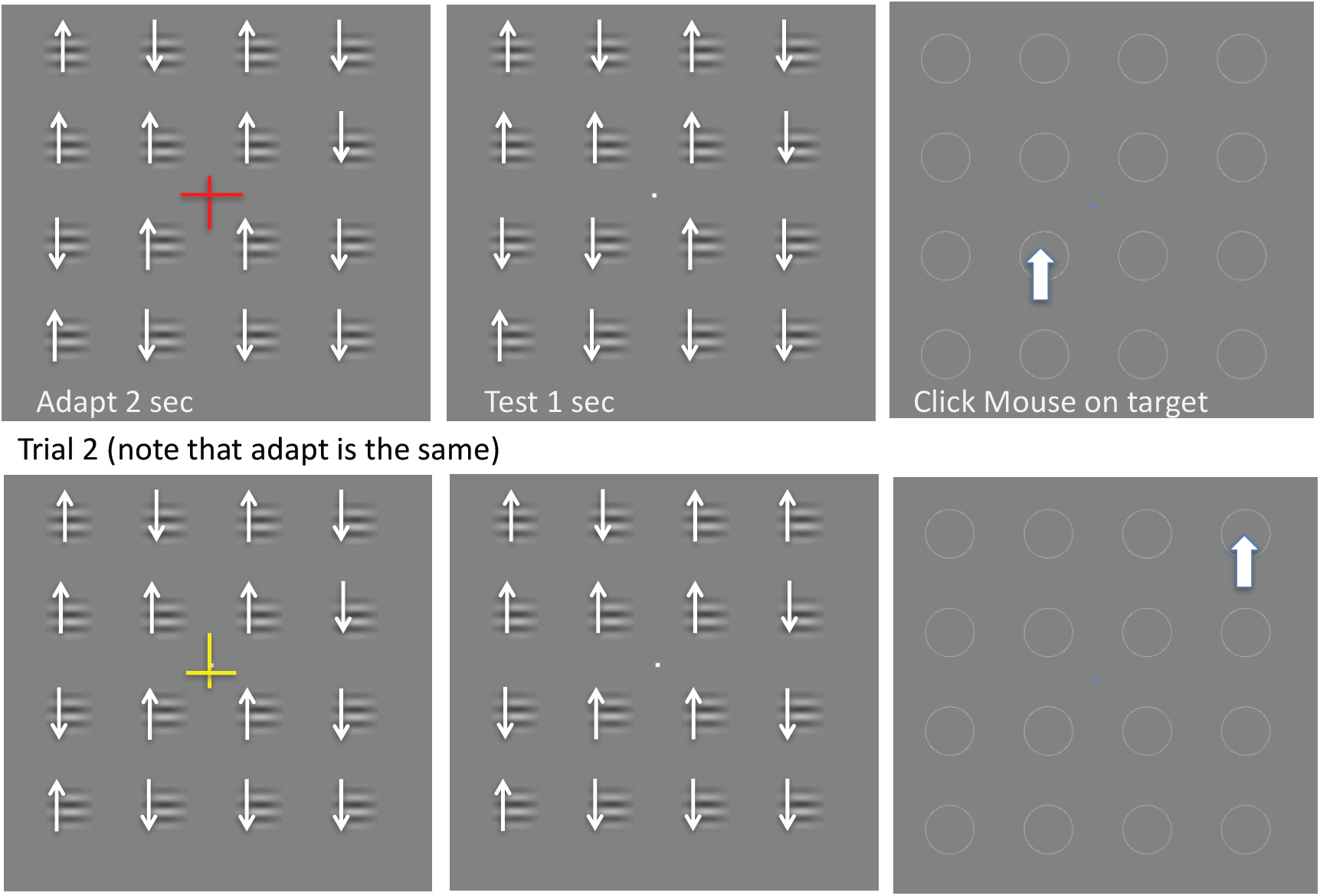
The figure shows a representation of two successive trials from Experiment 1. The arrows symbolize the direction of motion of each patch and were not visible during the experiment. Observers adapted to the array shown the top left panel while inspecting coloured crosses appearing at fixation at a rate of 1.5 Hz, and looking for rare combinations of shape (upright vs inverted) and colour (red, green, blue, or yellow). Here, an upright, red cross is illustrated. Each 2-s adapt period is followed by a 1-s test in which all the patches move in the same direction as their respective adaptors except the target, which moves in the opposite direction. The test is followed by presentation of circular placeholders, the observer’s task being to click on the position of the target. The placeholders remain visible until the mouse is clicked. In the experiment illustrated here the adapt and test arrays contained the same number of elements and the Gabor patches were all horizontal.

## General Method

### Apparatus and Subjects

Stimuli were presented on a 60-Hz frame-rate Sony Trinitron monitor in a darkened room, viewed from 0.75 m, so that one pixel subtended 1.275 arcmin at the observer’s eye. Viewing was binocular through natural pupils, with observers wearing their normal correcting lens for the viewing distance if necessary. A total of 12 observers participated in the experiments, comprising the three authors and a number of postgraduates/undergraduate students from City, University of London and the Max-Planck Institute for Metabolism Research at Cologne, all of whom were naïve to the purpose of the experiment. Two of the participants were paid volunteers.

### Stimuli

The stimuli (e.g. Fig. 1) consisted of rectangular arrays of Gabor patches, each of which comprised a sinusoidal grating multiplied by a circular Gaussian envelope. The grating had spatial frequency 3.75 cyc/deg drifting at 7.5 Hz and the Gaussian envelope had a spread (*σ* of 0.21 deg. The mean luminance and contrasts of the Gabors were 70 cd/m^2^ and 0.6 (60%) respectively. The envelope did not move. In most of the experiments it was truncated at ±2*σ*, so that it had a just-noticeable edge. In Experiment 2 truncation was at ±3*σ* Unless otherwise stated the array comprised 4 × 4 equally spaced Gabor patches, with a centre-to-centre spacing of 1.87 deg (8.75 *σ*).

### Procedure

Adaptation was produced by presenting one of these Gabor arrays for an initial 2, 3 or 5 s (in different experiments), during which the observer was instructed to fixate a stationary point in the centre of the display, and to carry out a task based on additional stimuli presented there. In Experiments 1 and 3, all Gabors were horizontal. Half moved upwards and the other half downwards. The spatial distribution was randomized but held constant within each session (see Fig. 1). In Experiment 2 Gabors could have any orientation (and thus move in any direction; e.g. Fig. 6).Nonetheless, their spatial distribution was always held constant within each session.The first adaptation period was followed, after 0.3 or 1 s (in different experiments), by a test, the duration of which varied in different experiments between 0.25 and 2 s.

With drifting adapters, change was introduced by reversing the direction of drift. With flickering adapters, it was introduced by rotating the carrier grating. After the test, the stimuli were replaced by a set of circular placeholders, and the observer used a mouse to click on the position of the target. To give feedback, the target’s placeholder was switched off to show the target’s position after the mouse click.

The first adapt-test cycle was followed after a blank interval of 0.5 sec, during which the screen was grey. It was then followed by other cycles of the same kind, in which the top-up adaptation period was 2, 5, or 15 s (in different experiments). The condition (Target-change or Distractors-change) was constant within a session.

All observers were given several sessions of practice before the main experiment in order to accustom them to the task, the aim being to start the main experiment only when they were detecting the target at above chance levels. However, this was not possible in observer EL who failed to show convincing evidence of detection. Two other subjects (DP, TP) also found he task difficult. EL, DP, and TP were included in the main experiments anyway, to see if they would eventually learn.

### The Central Attentional Distracting Task

To take attention away from the adapting stimulus during adaptation, observers in some conditions carried out a demanding RSVP task based on stimuli appearing at fixation. In the centre of the adapting array, superimposed on the white fixation point, a series of asymmetrical, coloured crosses were presented at a frequency of 1.5 Hz (2 crosses per trial in Experiment 1; 6 per trial in Experiment 2, and 3 per trial in Experiment 3). In the high-load (i.e. attentionally demanding) version of the task (see Morgan, 2011; Schwartz, Vuilleumier, *et al.*, 2005), the observer’s task was to press a button when there was a rare conjunction of colour and orientation. On all but 9.75% of the trials in Experiments 1 & 3 and 4.94% of the trials in Experiment 2, upright crosses were either red or green, and inverted crosses were yellow or blue. The four combinations were equally probable. On the remaining trials the first and following crosses were exceptions to this rule, for example the cross was red and inverted. Observers were instructed to press a key as soon as they saw an exception. As soon as they did so, the rule was reinstated. The observer was told that the exceptions were rare and that they should not produce false-positives. In the low-load versions of the task, the observer was instructed merely to maintain fixation on the crosses, or the crosses were absent.

## Experiment 1

### Methods

The stimulus configuration is shown in Fig. 1. The adapting duration, blank interval between adaptation and test, and the test interval were 3 s, 1 s, and 1 s. Each session began with 5 trials in which the adapting and test stimuli were absent, to accustom the observer to using the mouse to click on a placeholder, and to the appearance of the crosses. These initial trials were followed by 32 trials, with a total of 2 targets in each of the 16 positions, randomly interleaved without replacement. There were three attentional load conditions during adaption: crosses absent, high load, and low load (Conditions 1-3 respectively). In the low-load condition the crosses were present, but the observer had no task to perform. The colors were red, green, yellow and blue. All observers carried out at least 2 blocks of 32 trials each in each of the 3 Attentional Load conditions (6 blocks; 192 trials). Some observers did more. The actual total number of blocks performed by each of the observers (in order of their appearance in Fig. 2) was as follows: {25, 9, 13, 7, 6, 11, 18, 9, 6, 52, 10}. A total of 12 observers took part, but one of them (LP) did only the low-load conditions and her results are not presented in this section, although they are included in Fig. 9.

**Fig. 2.**
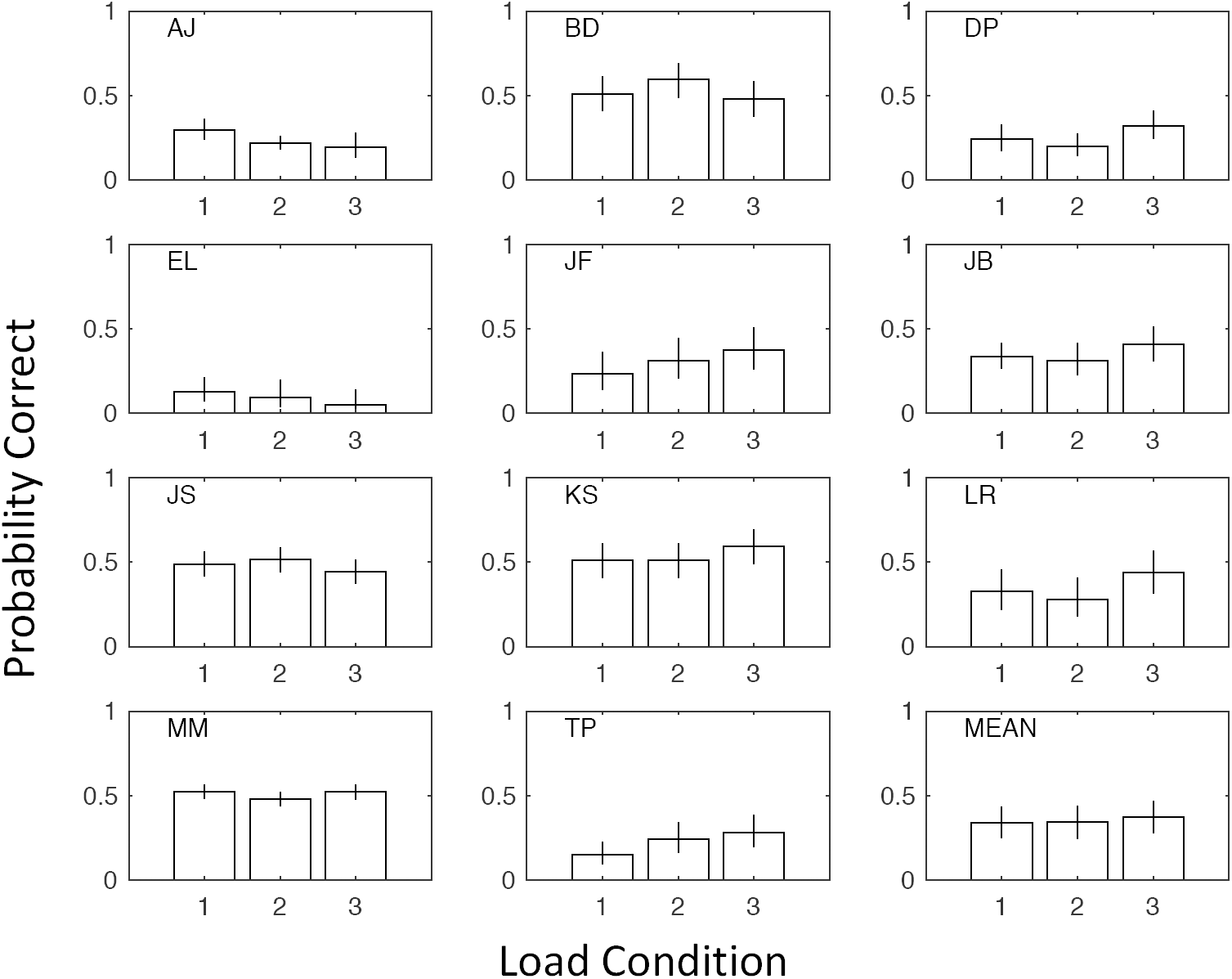
Performances of all 11 observers who completed Experiment 1 under three conditions of attentional load during adaptation (1: crosses absent, 2: high load, 3: low load). Observer LP is excluded from this analysis because she completed only the low-load condition. Error bars for individual observers contain binomial 95% confidence intervals. The bottom right panel shows the mean performance ± 1 SE over observers.

### Results

Fig. 2 shows that there was no systematic effect of attentional load during adaption on the overall success rate combined over trials and sessions, such as might arise from differences either in growth rate or asymptote of performance. To see specifically if load affected the build-up of adaptation during a session (as suggested by Bartlett *et al.*, 2016), growth curves over 8 successive 4-trial blocks (combined over sessions) were fit with the two-parameter function

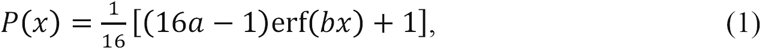

where *p*(*x*) is the probability correct on trial *x*, and parameters *a* and *b* describe the asymptote of performance and the growth rate, respectively (1/16 ≤ *a* ≤ 1, 0 ≤ *b*). The fits of the growth curves are shown in Fig. 3.

**Fig. 3.**
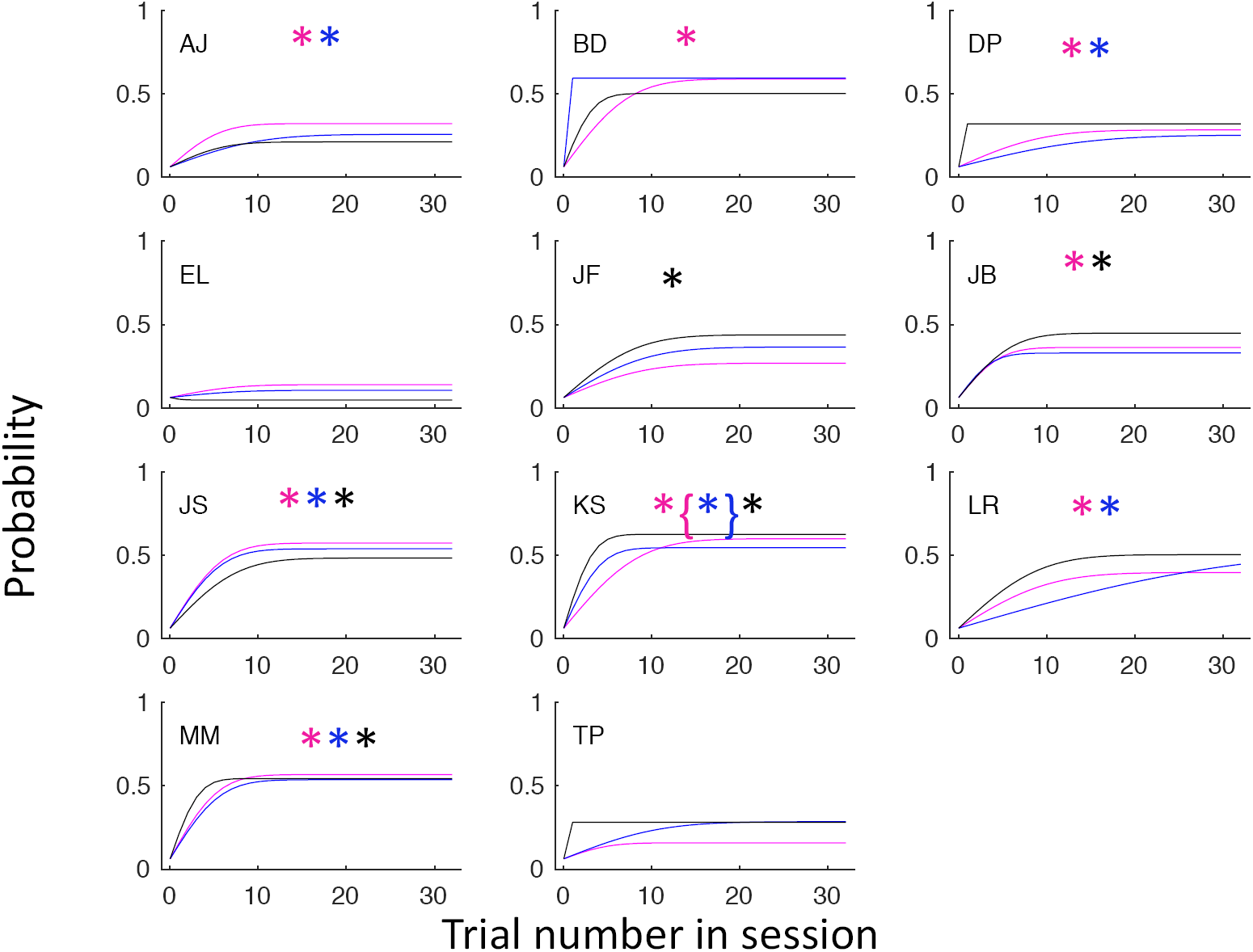
Growth curves in performance in Experiment 1 fit with the 2-parameter Equation (1) in the text. Each panel shows results for a different observer. The three curves are for the different conditions (magenta: crosses absent, blue: high load, black: low load). The colored symbols next to the observer’s initials indicate conditions in which the growth curve was significantly [χ2(1) > 3.84; p < 0.05] shallower than a step function (i.e. parameter *b* << 10); brackets indicate [χ2(1) > 5.64; p < 0.01]

Data from each observer were fit with Equation (1). Generalized likelihood-ratio tests (Mood, Graybill, & Boes, 1974) indicate that the fits did not improve significantly [in all cases *χ*^2^(2) < 5.99; p > 0.05] when parameters *a* and *b* were allowed to vary between conditions relative to the case where only *a* was allowed to vary between conditions, or relative to the case where only *b* was allowed to vary between conditions. At the group level (i.e. when responses were pooled across the first 6 sessions for each observer), fits did not improve when either parameter was allowed to vary over load conditions [*χ*^2^(2) < 0.18 ; p > 0.05]. This did not change when the three most poorly performing observers (EL, DP, TP) were excluded [Group Level *χ*^2^(2) < 0.95; p > 0.05]. We conclude that any effect of attentional load is small and inconsistent over observers, as is clear from inspection of both Figs. 2 and 3.

As a simple test of improvement during a block of trials, we used the generalized likelihood ratio test to compare the fit of Equation (1) when parameter b was free to vary with a fit with parameter b fixed at 10 (producing asymptotic performance for all trials, i.e. × ≥ 1). Using this criterion [*χ*^2^(1) > 3.84; p < 0.05], significant improvement occurred in the cases indicated by colored asterisks on Figure 3. At the group level, combining data over the same number of trials (64) for each obsever, the improvement was significant in all three conditions [*χ*^2^(1)> 7.25 ; p < 0.01].

An even simpler test is to subtract the mean success rate on trials 2–32 from the first-trial success rate. At the group level this difference score was negative (indicating improvement) in all conditions (−0.30, −0.28, and −0.20) for Conditions 1&An even simpler test is to subtract the mean success rate on trials 2–32 from the first-trial success rate. At the group level this difference score was nega#x2013;3, respectively). It was also negative for all observers in Condition 1, 10 of 11 obference due to perceptual load. Of course, the improvement could be general rather than due to adaptation, although all observers had been given several sessions of practice before the main experiment in order to accustom themselves to the task. To check this point, we analyzed performance in only the second block of 32 trials. This did not alter the situation: difference scores remained negative at the group level (-0.23, -0.31, and -0.37), for Conditions 1–3, respectively), and the numbers of obsthis point, we analyzed performance in only the second block of 32 trials. This did not alter the situation: difference scores remained negative at the group level (-0.2ervers with negative scores were 11, 10, and 10. We conclude that performanace improve

## Experiment 2

A possible criticism of Experiment 1 is that the adaptation periods were kept deliberately brief (2 s) in order to follow the build-up of adaptation over trials (32 per session). The periods may have been too brief to allow full deployment of selective attention to the crosses. Against this, we shall show later that detection rate for the exceptions (see General Methods) was high. However, as an additional precaution, in Experiment 2 the adaptation periods were lengthened to 5 s (6 crosses per trial). The high load condition was the same as in Experiment 1, and the low load was the same as Condition 3, i.e. crosses were present but required no response. The initial 5 trials of pre-adaptation in Experiment 1 were not used, as the observers were well used to the procedure. A further change from Experiment 1 is that the adapting array consisted of 6 × 6 elements of which only the central 4 × 4 were used for the test. This was in anticipation of a later experiment (not described here) in which the observers moved their eyes between adapt and test to demonstrate that the adaptation is retinotopic. The participants were the same as in Experiment 1, with the exception of LR and LP who did not take part.

### Methods

The general arrangement of the display is shown in Fig. 4, taken from screen-shots of the stimuli. The orientations of the patches were randomised between sessions, but constant within each 32-trial session. In anticipation of Experiment 4, we made the (6 × 6) adapting array larger than the (4 × 4) test array. We also truncated each Gabor’s Gaussian envelope at ±3*σ* to give it a softer edge than those used in Experiment 1.The initial and top-up adaptation periods were 5 s. These were followed by a 1-s blank, a 1-s test, and finally the 4 × 4 array of circular placeholders in the positions of the test patches. One of the moving patches changed direction through 180 deg between the adaptor and the test. The other patches continued in the same direction. Ten observers participated in the experiment, including authors MM and JS. The others were a mixture of postgraduates, colleagues, and paid subjects, all of whom were naïve as to the purpose of the study. One observer (AJ) was available for only 4 sessions (2 of each Load Condition). The others, in order of their appearance in Fig. 5 performed {12, 11, 12, 11, 7, 10, 14, 17, 17} sessions.

**Fig. 4.**
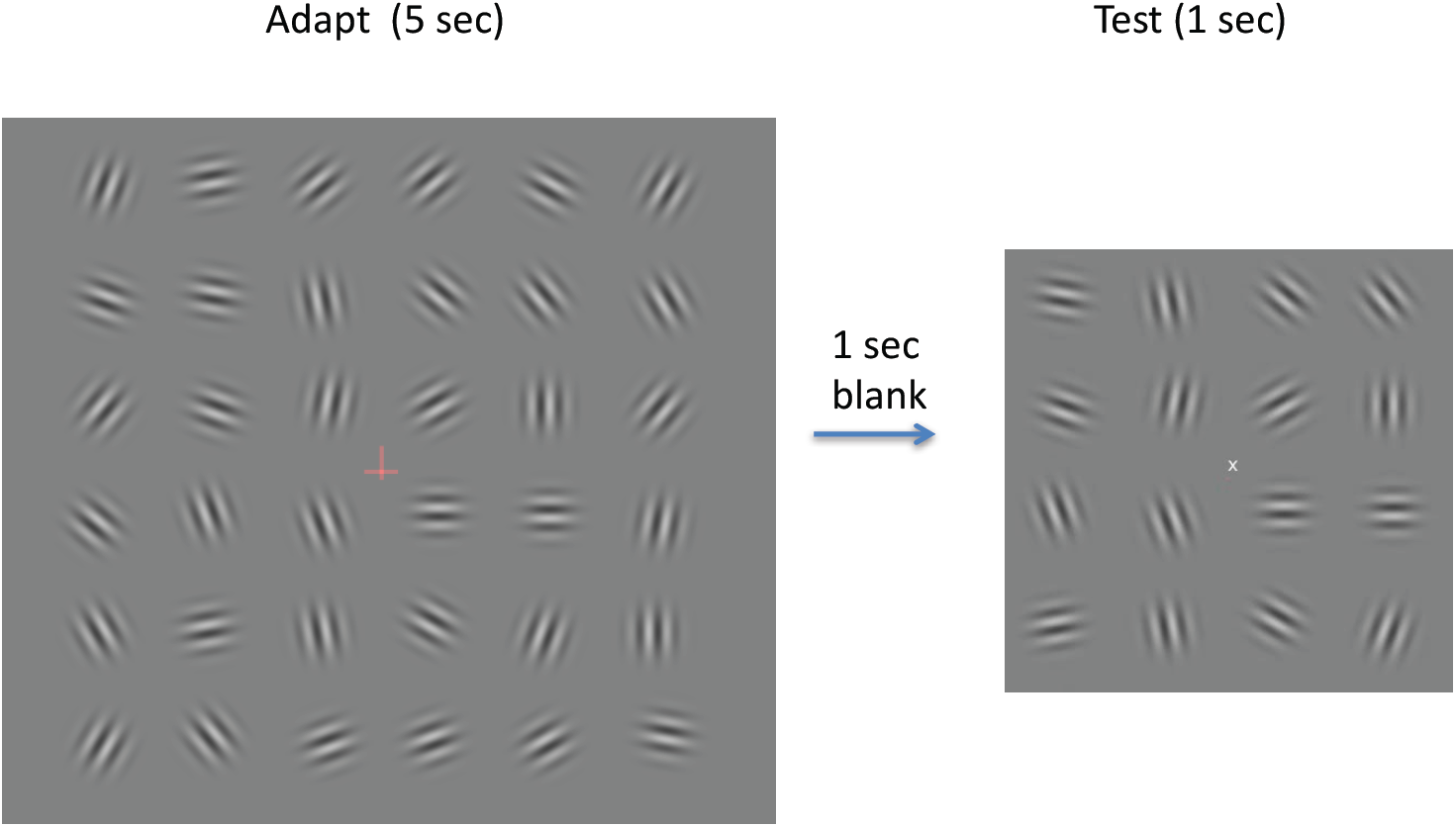
Stimuli used in Experiment 4. The central cross changed color and orientation 6 times during each adapting exposure.

**Fig. 5.**
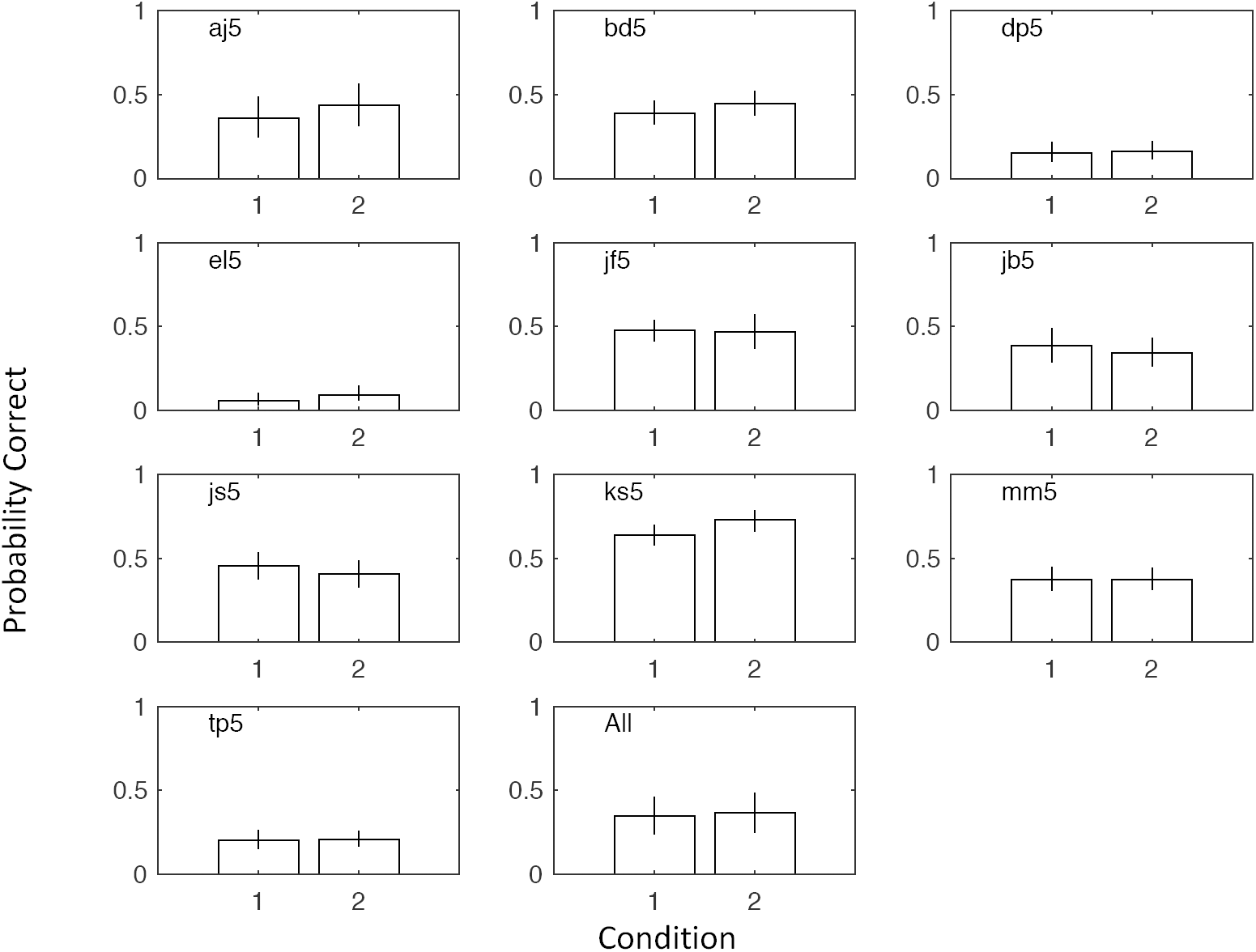
Performance of observers in Experiment 2 under two conditions of attentional load during adaptation (1: high load, 2: low load). Crosses were present in both conditions. Error bars for individual observers contain binomial 95% confidence intervals. The bottom right panel shows the mean performance ± 1 SE over observers.

### Results

The results summarised in Fig. 5 show that most observers performed the task with an accuracy between 0.4 and 0.6. Growth curves (Fig. 6) from each observer were fit with Equation (1). The only significant improvement [*χ*^2^(1) > 3.84; p < 0.05] when either parameter *a* was allowed to vary between attentional load conditions or when parameter *b* was allowed to vary between conditions, was found for slope in Observer KS [*χ*^2^(1) = 5.55; p < 0.05]. Nor did fits improve significantly at the group level, when either parameter was allowed to vary with load [in both cases *χ*^2^(1) < 3.84; p > 0.05].

**Figure 6.**
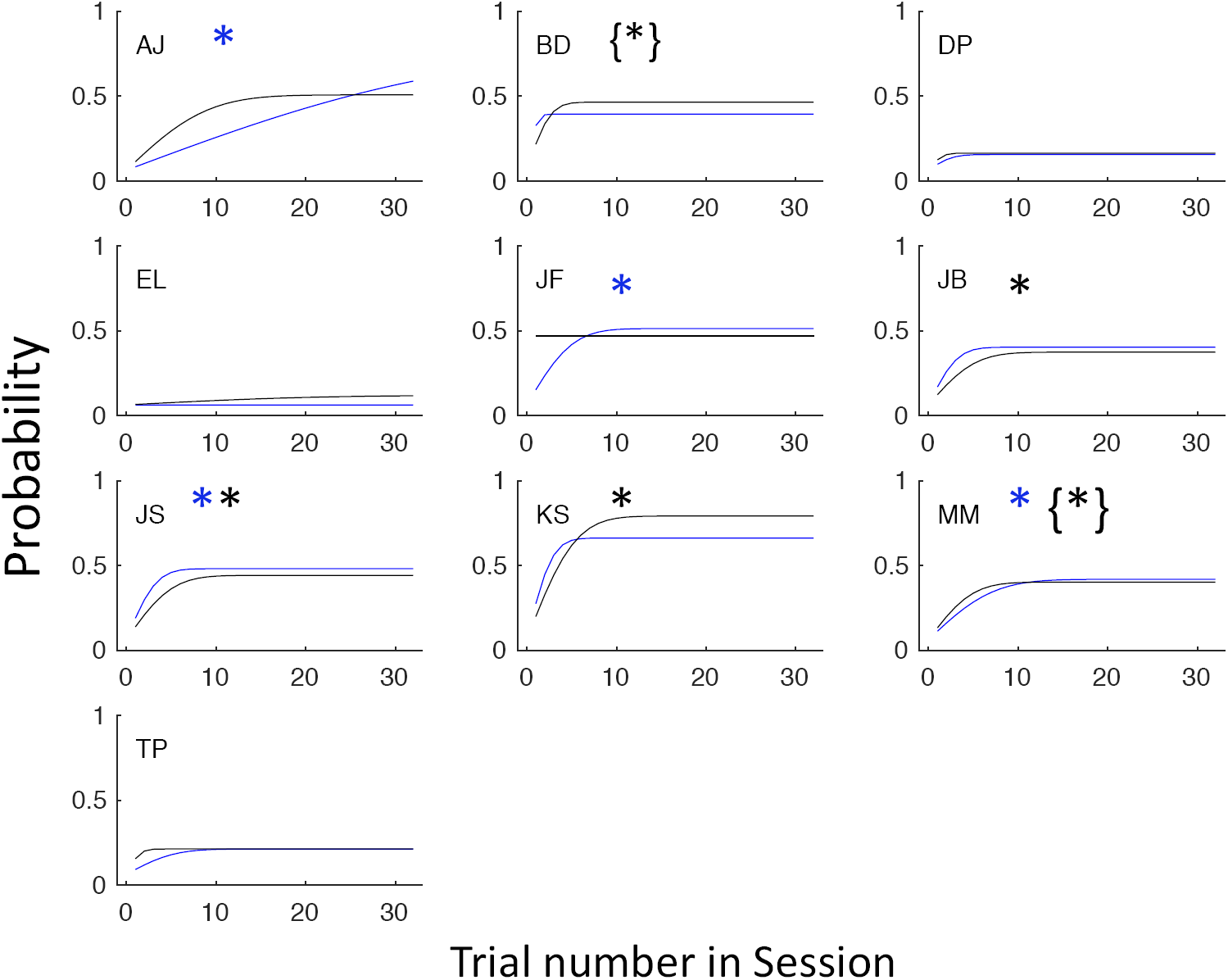
Growth curves in performance in Experiment 2 fit with the 2-parameter Equation (1) in the text. Each panel shows results for a different observer. The two curves are for the different attentional load conditions (blue: high load, black: low load). The coloured symbols next to the observer’s initials indicate conditions in which the growth curve was significantly (χ2(1) > 3.84; p < 0.05) shallower than a step function (i.e. parameter *b* << 10); brackets indicate [χ2(1) > 5.64; p < 0.01].

To test for a gradual build-up in adaptation over trials, we used the generalized likelihood ratio test to compare the fit of Equation (1) when parameter b was free to vary with a fit with parameter b fixed at 10 (producing asymptotic performance for all trials, i.e. × ≥ 1). Using this criterion [*χ*^2^(1) > 3.84 ; p < 0.05], significant improvement are indicated by colored asterisks on Figure 6. At the group level, combining data over the same number of trials (64) for each obsever, improvement was significant in the high load condition [*χ*^2^(1) = 6.62; p < 0.025] but not in the low load condition [*χ*^2^(1) = 1.85; p > 0.05]. The overall evidence for growth is thus not very strong. This may be because each adaptation period lasted for 5 s rather than the 2 s in Experiment 1, so that most of the build-up may have occurred on the first trial. To test this we carried out the same test as in Expt 1, taking a difference score between the first trial in each block and the mean of the remaining 31 trials. Although the difference scores at the group level were negative in both load conditions (-0.19 and -0.14) and was also negative for all observers in the high load condition, it was negative for just 6 of 10 observers in the low load condition. (NB: Just by chance, we would expect it to be negative in 5 of 10 observers.) A similar pattern was found when only the second block of trials was analyzed (group-level difference scores: -0.26 and - 0.07, number of obervers with negative scores: 9 of 10 and 7 of 10).Three of the observers (DP, TP, and EL) found the task very difficult and reported seeing no pop-out. Their scores barely reached the 0.25 level. DP and TP are siblings, and had performed normally in other psychophysical experiments, including motion direction discrimination (cf. Morgan, Schreiber & Solomon, 2017, where DP goes by the initials DW). Before concluding that these observers had reduced levels of adaptation, we wished to design an inverse task, where such a deficit would make the observer better, not worse. This was the purpose of Experiment 3.

## Experiment 3

In the Ishihara pseudo-isochromatic tests for colour deficiency, one of the plates contains numerals that should be invisible to normal trichromats but visible to dichromats (cf. Morgan, Adam & Mollon, 1992). We wished to develop an analogous test of adaptation, capable of identifying individuals whose efforts might be allocated more in accordance with their expectation (e.g. high load impairs search) than with task demands.

Our solution to this problem was suggested by the observation in Experiments 1 and 2 that the stationary, circular placeholders following the adaptation and test appeared to move, because of the well-known waterfall illusion (a. k. a. the motion aftereffect or MAE). We reasoned that this apparent movement of all the placeholders would camouflage real movement of one of them (analogous to Morgan, Adam & Mollon, 1992 for textural camouflage). Therefore, individuals with genuinely weak or absent adaptation should perform better than “strong adapters” in detecting the moving target. Amongst observers whose adaptabilities genuinely differ, we predicted a negative correlation between performances in the current experiment and those in Experiments 1 & 2.

### Methods

The adapting array and placeholder array were the same as in Experiment 2, but the test array was omitted. The adapting duration was 3 s, followed by a 1-s blank screen, and finally the placeholder array, which remained on until the observer clicked on the moving target. The measure of performance was the reaction time (RT) before the click; shorter RTs indicate better performances. During the display one of the circular placeholders moved vertically at a slow speed of 8.75 arcmin/s, either in the same direction as its spatially corresponding adaptor or in the opposite direction. The target-same and target-opposite conditions were randomly interleaved within a block, with an equal number of each. Because all the placeholders seemed to move initially, the observer had to wait for the MAE to die down before deciding which one was really moving. Typically, this took a few seconds. In a control condition, the patches in the adapting array were stationary. In this case, the distinction between target-same and target-opposite conditions was purely notional.

Attentional load was manipulated with central crosses, as in Experiment 2. In the high-load task there were 3 crosses in each adapting interval. In the low-load condition there were no crosses, as in Condition 1 of Experiment 1. The initial 5 trials of pre-adaptation in Experiment 1 were not used, as the observers were well used to the procedure. The observers were the same as in Experiment 1

### Results

A simple summary of the individual RTs in Fig. 7 is that the all the conditions are equivalent, except for the case where the adaptor is moving and the target is moving in the same direction as the adaptor. Detection was slower for targets moving in the same direction (T. same) as its corresponding adaptor than for targets moving in the opposite direction (T opp.). Of course, this difference was not found in the case where the adaptor was static (Stat.); only when it was moving (Mov.). The asymmetry is easily explained if adaptation to a moving Gabor patch causes a perceived movement of the spatially corresponding placeholder circle in the opposite direction. If the real movement is in the same direction as the adaptor, the aftereffect will slow it down and make the movement harder to detect. If it is in the opposite direction, the aftereffect and real motion will add and make it easier to detect. Note, however, there seem to be genuine individual differences. Observers EL, DP, and TP do not show the predicted longer reaction times in the target-same condition. These are the same three observers who found the task particularly difficult in Experiment 2, where pop-out was based on prior adaptation. We therefore conjecture that these observers are “weak adapters.”

**Fig. 7.**
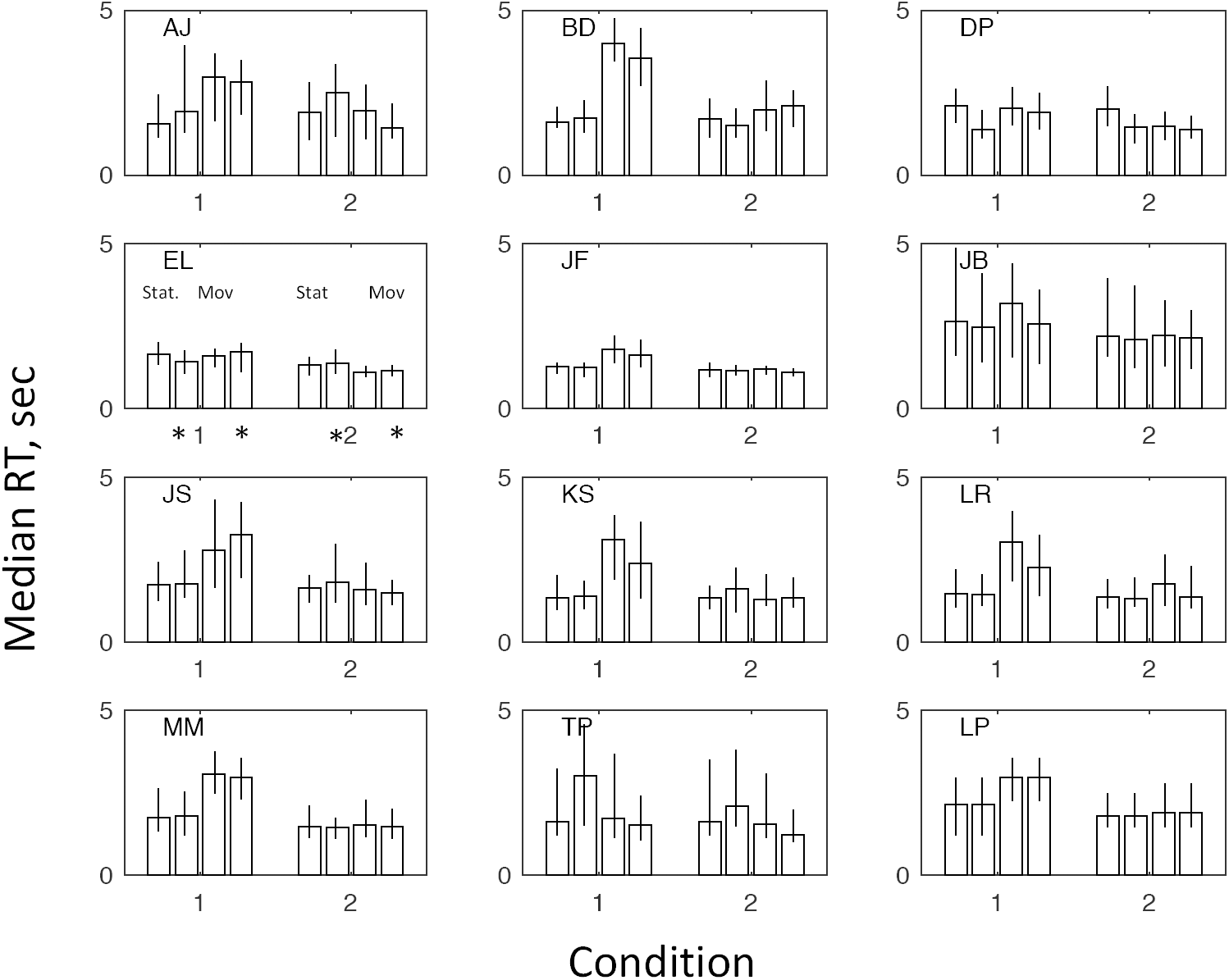
The figure shows the data from Experiment 3, in which observers identified a slowly moving target circle amongst 15 stationary distractors, as quickly as possible, after adapting to a 4 × 4 array of Gabor patches (see Fig. 1). The key to the conditions is given in Panel EL. The adapting patches were either static or moving (Stat. vs Mov.). In the adaptor-moving conditions, the target moved either in the same direction as its preceding adapting patch (Condition 1) or in the opposite direction (Condition 2). In the adaptor-static conditions, the target was randomly assigned to the target-same or target-opposite condition. The * symbol indicate that the adaptation was carried out under a high attentional load condition. Observer LP carried out the low load condition only; her data for this condition have been repeated. The vertical axes show the individual median times taken by the observer to click on the target. The error bars contain 50% of the data.

The error rate in target selection was low (grand mean over all observers and conditions: 0.088) and not significantly different between conditions. Observer AJ was unusual in having a relatively high error rate (0.2891).

The effects of attentional load were not consistent over observers. However, if we set aside the results for the three weak adapters, we find that 7 out of the 8 remaining observers show longer reaction times in the low-load condition, consistent with the combined data (bottom right panel in Fig. 8). The exception is JS. However, it must be acknowledged that the effect of load is also miniscule in observers MM and AJ. Overall, the effect of load is not very convincing.

**Fig. 8.**
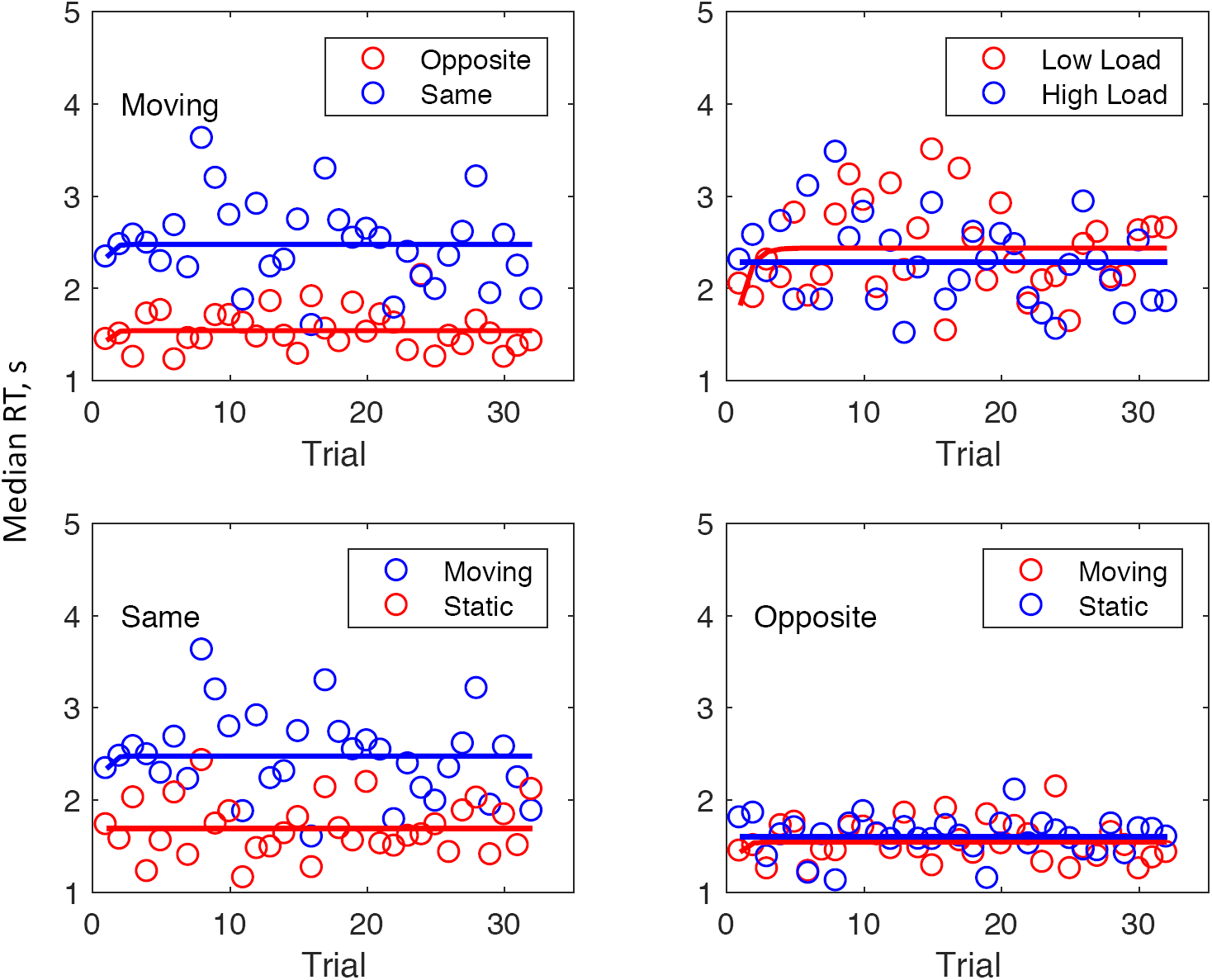
Each panel in the figure shows the median reaction times over observers (ordinate) plotted against trial number in the session (abscissa). The solid lines are fits to the data points of the same colour using the negative exponential growth curve described in the Text. Each panel shows a different set of conditions, as follows: *Top Left*: Contrasts targets moving in the same direction as their adaptor with targets moving in the opposite direction. Only trials with a moving adaptor are included. Load conditions are combined. *Top Right*: Contrasts the two conditions of attentional load. Only moving adaptor + target-same conditions are included. v*Bottom Left*: Contrasts the moving-adaptor and static-adaptor conditions, when the target moves in the same direction as the adaptor. Load conditions are combined.*Bottom Right*: Contrasts the moving-adaptor and static-adaptor conditions, when the target moves in the opposite direction from the adaptor. Load conditions are combined.

To investigate possible effects of attentional load on the build-up of adaptation we first fitted negative exponential growth curves to the median RTs over observer (Fig. 8). Surprisingly, in view of the results of the first two experiments, there was little evidence for an increase in RT, as would be expected from build-up of adaptation, even in the crucial condition where the test and adapting stimulus moved in the same direction (Panel 1 in Fig. 8), except possibly from Trial 1 to Trial 2. It was not possible to test for growth using the same likelihood tests used in Experiments 1 and 2 because the data were RTs rather than probabilities, so we used the extra-sum of squares F-test (Mood, Graybill & Boes, 1974):

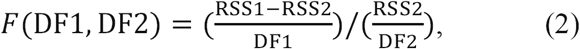

where RSS2 is the residual sum of squares after fitting with the more complicated model with two free parameters (an asymptote *a* and slope *s*), and RSS1 is for the nested model, with the slope fixed at *s* = 10 and a single free parameter (*a*). The degrees of freedom are DF1 = 1, the difference in free parameters, and DF2 = 30, the difference between the number of data points (32 in this case) and the the number of free parameters in the more complicated model. Tests on individual observers in the crucial condition where we expected adaptation (see above) revealed no case, in either condition, where the nested model fit significantly worse than the more complicated model [in all cases *F*(1, 30) < 2.19, where the critical value for *p* < 0.05 is 4.11). At the group level, testing the median RTs over observers, no *F* was greater than 1.006 (NS). The difference scores between the first trial and the median of trials 2-32 were negative in 6 of 10 and 4 of 10 observers in the high and low load conditions, respectively (group data shown in top right panel of Fig. 8). We conclude that the evidence for growth of adaptation is weak, despite the fact that adaptation clearly had an effect (as shown in the top left panel of Fig. 8, for example).

We next used the same F-test to test the significance of the differences in asymptote (or slope) between the pairs of conditions in each of the 4 panels of Fig. 8. At the group level, using medians over observers as data, the differences in asymptote were significant for the effect of movement direction [top left panel in Fig. 8; *F*(1, 30) = 40.6, *p* < 0.001] and for the effect of moving vs static adaptor [*F*(1, 30) = 30.66, *p* < 0.001] but not for the effect of load [top right panel *F*(1, 30) = 1.08, *p* > 0.05] nor for the contrast in the lower right panel [*F*(1, 30) = 0.06, *p* > 0.05]. None of the differences in slope between pairs of conditions in Fig. 8 were significant (*F* < 0.36 in all cases). These statistics confirm the visual impressions from Fig. 8, and the results of analyzing individual observers, for none of whom was the effect of load significant (all *F*s < 1.53; *p* > 0.05).

To investigate further the differences between observers, we assigned each observer a score in Experiment 1 (the detection probability, or hit rate) and plotted this against the difference score between median reaction times in the target-same and control (i.e. static adaptors, in the notional ‘target-same’ condition) conditions in the present experiment (data from both high and low-load conditions were pooled). The difference score on trial *t* was computed as (RT1(*t*)-RT2(*t*))/(RT1(*t*)+RT2(*t*)), where *t* is the trial number and RT1(*t*) and RT2(*t*) are the individual reaction times on that trial in the two conditions respectively. The points Fig. 9 are the medians of the set of difference scores and the error bars contain 95% of the set over all trials. We expected low detection probabilities in Experiment 1 to correlate with low difference scores in Experiment 3, because both imply weak adaptation. The results shown in Fig. 9, comparing Experiments 1 and 3 confirm this conjecture. The three putatively weak adapters (EL, DP, and TP) form a distinct cluster in the bottom left of the figure. The overall correlation (ρ) is 0.67. A similar analysis of Experiments 1 & 2 showed the same clustering (Fig. 10) for 12 observers.

**Fig. 9.**
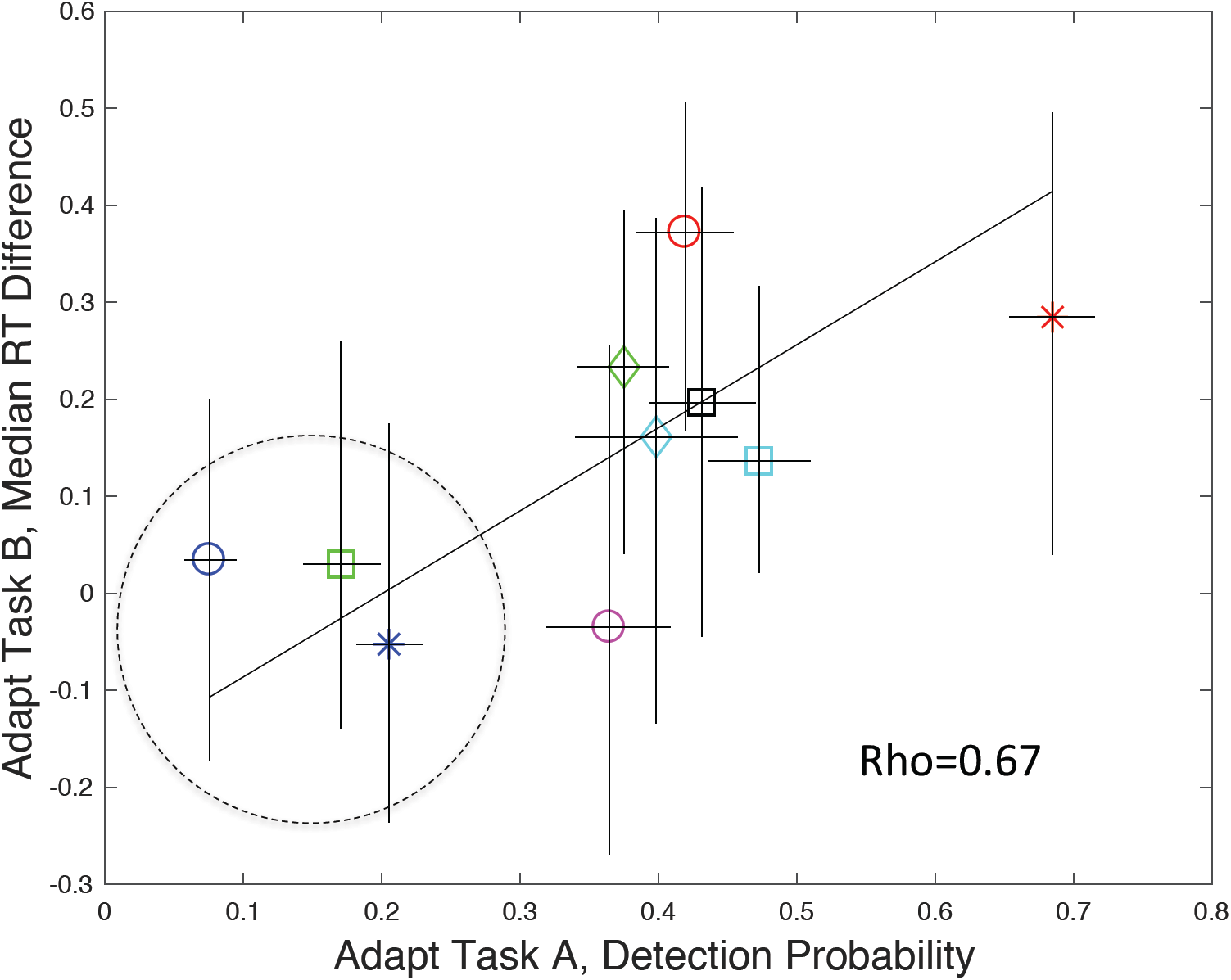
Median (±47.5 percentiles) difference scores for 10 observers in Experiment 3 (ordinate) plotted against hit rate for the same subjects in Experiment 2. Observers DP, EL, and TP (green square, blue circle, and blue star) form a distinct cluster, indicated by the dotted circle) in the bottom left of the space. Horizontal error bars contain binomial 95% confidence intervals. The straight line shows the best-fitting linear relationship between Task A probability and Task B RT difference. Data have been combined across Load Conditions in both experiments.

**Fig. 10.**
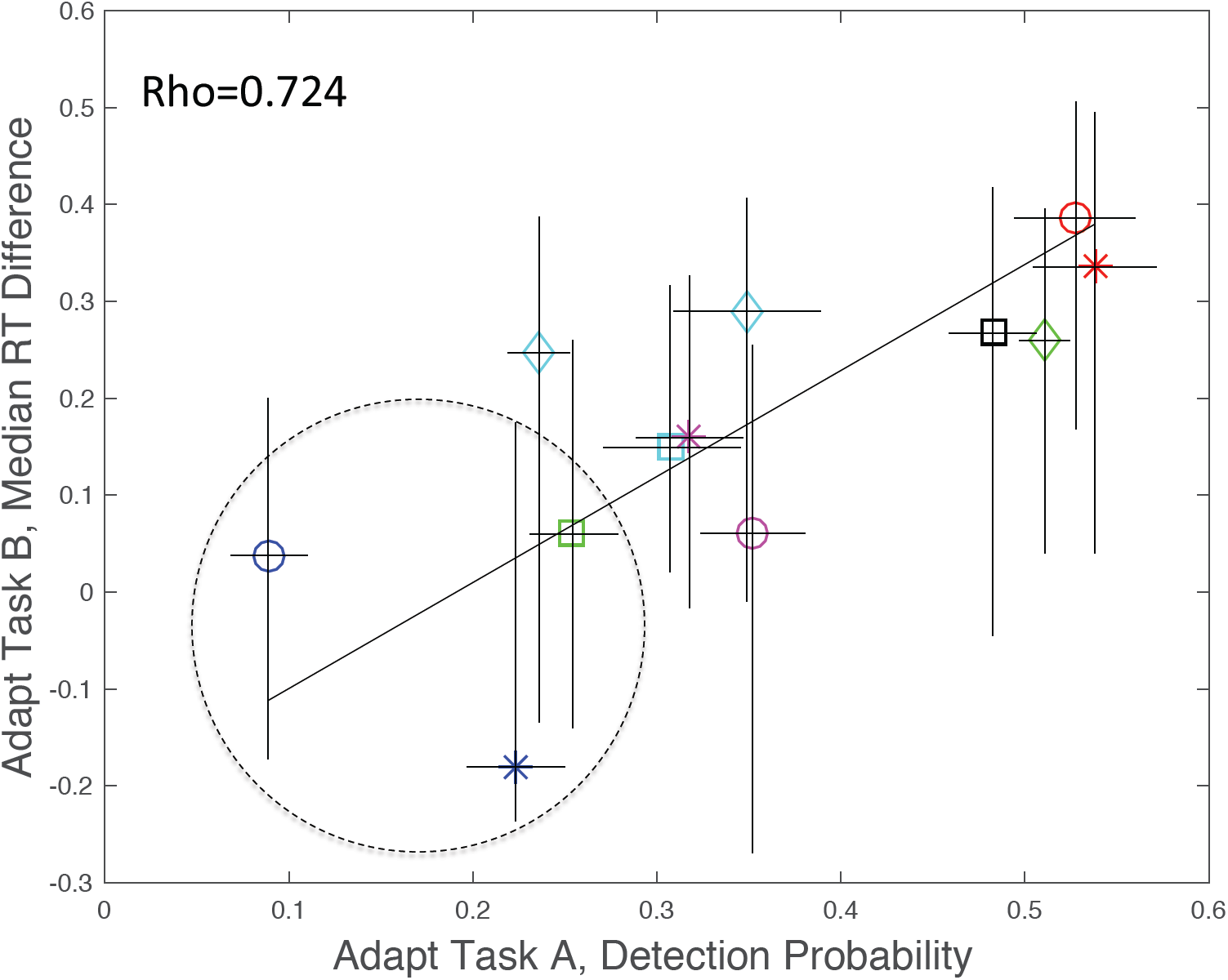
Difference scores for 12 observers in Experiment 3 plotted against hit rate for the same subjects in Experiment 1. Layout analogous to Fig. 9. Observers DP, EL, and TP (green square, blue circle, and blue star) again appear in the bottom left of the space. A data point for LP (magenta star) is included, although she experienced only low-load conditions in both experiments.

### The Central Distracting Task in Experiment 1-3

To test whether observers were actually paying attention to the crosses in the distracting task, we analysed their success rate and reaction times in spotting the rare exceptions. Due to an intermittent keyboard connection error, which was only discovered after the data had been collected, data were not available for all subjects in all sessions, but the remaining data were sufficient to show that detection was indeed occurring. The mean hit rates over observers and the mean RT are shown in Table 1, with the standard deviations in parentheses. The longer RTs and higher hit rates in Experiment 2 are most likely due to the fact that observers had longer to respond (5 s) than in Experiment 1 (2 s) or in Experiment 3 (3 s). To see if there was any relationship between perceptual load and hit rate, we calculated a summary statistic for the effects of load. In Experiments 1 and 2 this was the difference in target detection probability between high and low load conditions. In Experiment 3 it was the difference in RT between high and low loads in the condition where the target moved in the same direction as the adaptor. The Kendall Correlation coefficients between these three measures and the overall hit rates were 0.11, -0.09 and 0.14 respectively, giving no evidence for an association.

| Experiment | P(hit) | RT (s) |
| --- | --- | --- |
| 1 | 0.609 (0.117) | 1.39 (0.18) |
| 2 | 0.78 (0.22) | 2.7 (0.86) |
| 3 | 0.72 (0.15) | 1.79 (0.42) |

### Eye Movement Recording

Eye-movement recordings were carried out to verify that observers maintained fixation during the adaptation top-ups (Fig. 11). These recordings also showed that the three putatively weak adapters (EL, DP, and TP) were just as accurate at fixating during adaptation as the other observers.

**Fig. 11.**
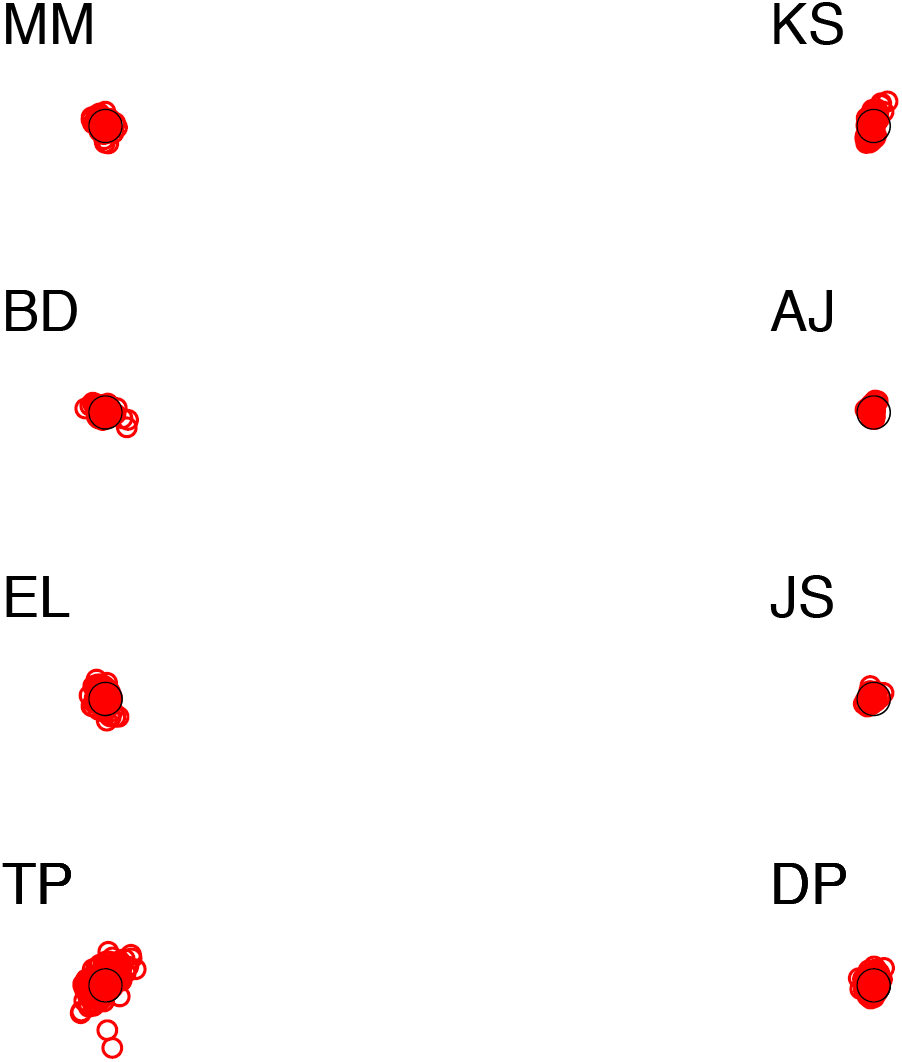
The data show mean gaze positions on separate trials, for 8 observers during adaptation top-ups (red) and during tests (blue). The centre circle is centred on the position of the FP during adaptation and is of similar size to the placeholders used in the experiments.

## Discussion

### Individual Differences

Exceptional individuals can help us to understand the physiological mechanism of sensory processing (Mollon, Bosten et al, 2017). Dalton’s anomolous colour vision, for example, greatly influenced the theory of trichomacy. However, for an exceptional individual or individuals to be informative, it is important that they be subject to rigorous testing; simply being poor in one particular test is not sufficient, because this could arise from many motivational and cognitive factors having little or nothing to do with the mechanisms of sensory processing. Particular care must be taken to disguise from the particants to purpose of the experiment and the result expected from the experimenter’s hypothesis (Manning, Morgan et al, 2017; Gheri,. Chopping & Morgan, 2008)

We found three observers who showed small or non-existent effects of adaptation in a visual search task. Crucially, this was not because they were poor at search tasks in general; they were faster than most observers in searching for the moving placeholder in Experiment 3, a task that was impeded by adaptation. Nor was it because these observers were relatively poor at fixating the centre of the adapting stimulus (Fig. 11). We conclude that these observers most likely have a weaker MAE than the norm. Large individual differences in the duration of the MAE have previously been reported by Granit (1928) and Sinha (1952), including the remarkable and neglected case noted by Grindley (1930) of an observer who saw no movement aftereffect whatsoever. The exciting possibility of a basic polymorphism in adaptation between human observers deserves further investigation. The advantage of the method we have described in this paper is that, unlike the duration or P50 measures of the MAE (Morgan, Dillenburger, Raphael, & Solomon, 2012), it is performance–based and criterion free. It is a “Type 1” task, for which there is a right answer (Sperling *et al.* 1990), and which therefore could be used to measure adaptation in non-human species.

The number of participants in our experiments (n=12), although large by the normal standards of psychophysical experiments, does not permit very accurate estimates for the prevalence of weak adapters within the university population, which may not be representative of the general population. Given our 3-out-of-12 incidence, the 95% binomial confidence interval extends from 0.07 to 0.57. Nor can we be confident that weak adapters form a distinct cluster, rather than being at one end of a continuous distribution. Nor can we say anything about the generality of the trait with respect to forms of adaptation other than the motion after-effect. To overcome these limitations, it would be necessary to run a larger scale study, with many more particpants and different kinds of adaption, such as orientation and contrast.

### The effect of attentional load

As noted in the Introduction, there are conflicting claims regarding whether the MAE is decreased when observers’ attention is distracted away form the adapting stimulus. Bartlett *et al.* (2016) recently suggested that the discrepency between these results might be because attention affects only the growth of adaptation to asymptote rather than the final level. Negative studies may have missed the growth effect. Specifically for this reason, we looked at the growth of the adaptation effect in Experiments 1 & 2. Within-subjects analyses of our objective (or “Type 1”) measure of adaptation produced no convincing evidence for differences in growth rate or asymptote. A small effect of load was found in Experiment 3, but only at the population level and when weak adapters were excluded. Contrast in this experiment was between crosses-present (high load) and crosses-absent (low load) so we cannot exclude the possibility that any small effect was due to the crosses rather than to attentional distraction *per se*.

It seems reasonable to conclude, taking into account the literature as a whole, and the negative effects reported in this paper, that the effect of attention on motion adaptation is at best small and inconsistent. Even if a recipe for producing the effect consistently were eventually found, it would be necessary to eliminate the possibility that it was due to peripheral effects of attention, such as microsaccade frequency, pupil size, and blinking, before we could conclude that it is a direct effect of attention on V1. Our view is that the effect, if it exists, is so small and variable over observers that it is not worth pursuing further in any detail.

## Acknowledgements

This work was supported in part by RPG-2016-124 from the Leverhulme Trust.

